# Concentration Sensing in Crowded Environments

**DOI:** 10.1101/2020.10.02.324129

**Authors:** Wylie Stroberg, Santiago Schnell

## Abstract

Signal transduction within crowded cellular compartments is essential for the physiological function of cells and organisms. While the accuracy with which receptors can probe the concentration of ligands has been thoroughly investigated in dilute systems, the effect of macromolecular crowding on the inference of concentration remains unknown. In this work we develop a novel algorithm to simulate reversible reactions between reacting Brownian particles. This facilitates the calculation of reaction rates and correlation times for ligand-receptor systems in the presence of macromolecular crowding. Using this method, we show that it is possible for crowding to increase the accuracy of estimated ligand concentration based on receptor occupancy. In particular, we find that crowding can enhance the effective association rates between small ligands and receptors to a large enough degree to overcome the increased chance of rebinding due to caging by crowding molecules. For larger ligands, crowding decreases the accuracy of the receptor’s estimate primarily by decreasing the microscopic association and dissociation rates.

**SIGNIFICANCE:** Developing an understanding of how cells effectively transmit signals within or between compartments under physical constraints is an important challenge for biophysics. This work investigates the effect that macromolecular crowding can have on the accuracy of a simple ligand-receptor signaling system. We show that the accuracy of an inferred ligand concentration based on the occupancy of the receptor can be enhanced by crowding under certain circumstances. Additionally, we develop a simulation algorithm that speeds the calculation of reaction rates in crowded environments and can be readily applied to other, more complex systems.

## INTRODUCTION

The ability of cells to gather information about their environment is essential for remaining viable in the face of changing surroundings. Of particular importance is chemoreception – the process through which cells measure chemical concentrations of their environment. In their seminal work on chemoreception, Berg and Purcell (1) determined a physical limit to the accuracy of concentration measurement due to the rate of diffusional encounters between reactants. This limit has since been extended to account for finite reversible reaction rates of ligand-receptor pairs in contact (2), as well as other chemoreception processes such as sensing by arrays of receptors (3), estimation of dynamical signals (4, 5), and measurements of concentration gradients (6). In these cases, it is generally assumed that the concentration of ligands is low and the system is dilute.

While this approximation is often valid for receptors located on the exterior surface of cells, many receptor systems operate within the cytoplasm or organelles such as the endoplasmic reticulum and nucleus. The cytoplasm and organelles differ from the exterior of the cell in that they are densely crowded with a diverse array of macromolecules. Still, receptors must be able to detect the abundance of molecules in order to activate feedback loops regulating cellular homeostasis. The inference of concentration is fundamentally dictated by the distribution of times for which the receptor is unoccupied (7), which in turn is dictated by a combination of the encounter rate and intrinsic reaction rate of the ligand and receptor. The accuracy of such an estimate increases as the number of independent measurements (i.e. binding events) increases. Since each binding event necessarily requires the previously bound ligand to have dissociated, the measurement error depends on the intrinsic dissociation rate of the ligand and receptor, along with the association and encounter rates (2). Macromolecular crowding influences each of these rates (8–10) suggesting it will affect the accuracy of sensing and concentration measurements within cells (11).

While signal transduction in the crowded cellular milieu is essential to many physiological processes, it has been studied far less thoroughly than signaling in dilute systems. A critical factor for cell signalling is the residence time of the ligand on the receptor. A longer residence time kinetically affects the activation of signaling pathways and is critical for drug development (12). The effects of macromolecular crowding on the ligand residence time remain largely unexplored. One approach to study the effect of crowding on intrinsic noise of a bimolecular reaction is to perturb the chemical master equations from the dilute limit (13). This relates the correlation time of the reaction to the intrinsic association and dissociation rates in a crowded environment. However, since the approach is based on the chemical master equations, the model does not account for spatial correlations nor diffusion explicitly. A full description of the signaling noise of a bimolecular reaction in a crowded environment requires a spatial description capable of capturing the structure of the fluid around the reacting molecules.

The importance of fluid structure is evidenced by simulations of irreversible reactions with macromolecular crowding. In particular, simulations of diffusion-limited reactions of dense hard-sphere fluids are reasonably well described with Smoluchowski theory (14) by correcting for changes to the radial distribution function and self-diffusion coefficient due to crowding (15). A similar approach, but with a finite association rate, showed that crowding could either increase or decrease the effective binding rate by slowing the rate at which distant ligand molecules approached the receptor while also increasing the amount of time ligands spend near the receptor once they arrive (16). These competing effects can lead to non-monotonic dependence of the association rate on crowding.

Reversible reactions differ from their irreversible counterparts due to the presence of high-frequency rebinding reactions and altered binding equilibria. Spatial correlation of these effects, along with slowed diffusion, have been explored in the context of enzyme catalyzed reactions (17), signal transduction (18), and transcription factor binding (19–22). In the latter, it is found that the effects of crowding can increase bursting in transcription by increasing the time spent by the transcription factor near the target gene. The ability of crowding to alter correlations in the binding of transcription factors and ligands suggests the accuracy of chemical concentration measurements based on reversible diffusion-influenced reactions should also be crowding-dependent (11). Yet, crowding’s impact on measurement error remains unexplored.

One hurdle to calculating the accuracy of a chemoreceptor from simulation is that long trajectories are required to gather robust statistics of binding events. This is exacerbated for crowded systems in two ways. First, crowding slows diffusion and, in some cases, lengthens the residence time (i.e. time a ligand will remain bound), thus increasing the length of trajectories needed. Second, crowding necessarily involves higher density systems which have many more pair interaction between particles. Hence, calculating reversible binding statistics in crowded environments can be computationally demanding.

To address this issue, we develop a novel approach to calculating survival probabilities and correlation functions for receptors in crowded environments. The method combines Brownian dynamics simulations with offline Monte Carlo sampling of reaction probabilities along trajectories. This allows for rapid simulation across a range of intrinsic rate constants. Using this method, we compare the accuracy of the chemoreceptors in crowded environments for small and large ligands. Interestingly, we show that under certain conditions, increased crowding can increase the accuracy of chemoreception. Additionally, we demonstrate how the residence time of the ligand on the receptor is heavily affected by the degree of macromolecular crowding. This result has important implications for drug development under physiological conditions.

## METHODS

### Sensing Error for a Binary Receptor

In this section we briefly review established results for the accuracy with which the concentration of a ligand species can be inferred from the fractional occupancy of a single receptor. Let the number of ligand molecules in a compartment be *n_L_* and be constant across the timescale over which the sensor makes a measurement, leading to a concentration *c_L_* = *n_L_V*^−1^ for a compartment of volume *V*. The receptor attempts to estimate the concentration *c_L_* based on the ratio of time spent bound to a ligand versus unbound, i.e. the fractional occupancy. In particular, the receptor can estimate the probability of a ligand being bound by averaging the occupancy, *n*(*t*), of the receptor over a time, *T*, such that the estimated occupancy is given by

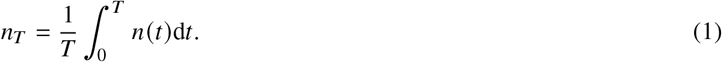

The estimate of the ligand concentration then follows from inverting the probability that the receptor is bound at a given concentration, *p*(*c_L_*), such that *c_L_* = *p*^−1^ (*n_T_*). The error in the concentration estimate, *δc_L_*, can be related to the variance in the estimate of the occupancy, 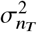, by (23)

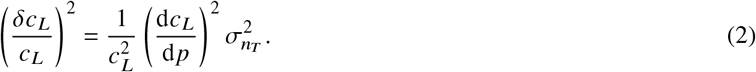

For the ligand-receptor binding considered here, the bound probability is *p*(*c*) = *c*/(*c* + *K_D_*) where *K_D_* is the ligand-receptor dissociation constant, and the gain of the sensor is *d*_p_/d*c*_*L*_ = *p*(1 – *p*)/*c*. Under the assumption that the measurement time is much greater than the autocorrelation time, *τ_C_*, of the receptor occupancy (i.e., *T* ≫ *τ_C_*), the variance in estimated occupancy is given by

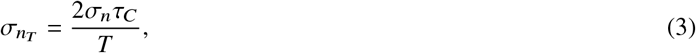

where 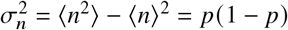 is the instantaneous variance of the signal. Substituting Eq. (3) into Eq. (2) leads to

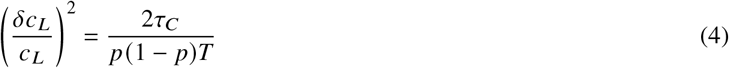

Hence, all that is needed to compute the sensing error is the autocorrelation time of the sensor occupancy.

The autocorrelation time is defined in terms of the autocorrelation function *C_n_* (*τ*) as

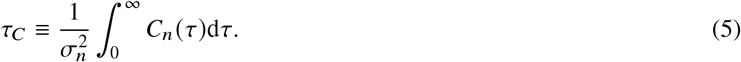

The correlation function for a binary switching process is given by

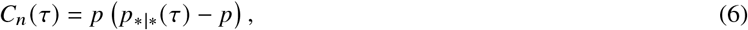

where *p* is the mean occupancy of the sensor and *p*_*|*_ (*τ*) is the probability that a receptor that was bound by a ligand at time *t* = 0 is also bound at time *t* = *τ*, allowing for potentially many binding and unbinding events between 0 and *τ*. In practice, it is easier to compute the complementary separation probability, 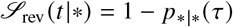, which gives the probability that a receptor which was bound at time *t* = 0 is unbound at time *t* = *τ*. It has been shown that the separation probability can be written as a convolution with the survival probability of a receptor with a ligand at contact (24)

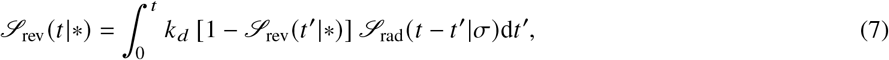

where *k_d_* is the microscopic dissociation rate of the bound ligand-receptor complex. 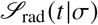 is the survival probability of a receptor from which a ligand dissociated at time *t* = 0 and *σ* represents the separation of the centers of the just-dissociated ligand-receptor pair. The subscript “rad” is used to indicate that the survival probability relates to the irreversible reaction case in which the receptor acts as a sink with a radiation boundary condition, as opposed to the subscript “rev”, which denotes the reversible reaction case. In general, a closed form expression for 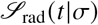 does not exist. However, under conditions in which the rebinding of the neighboring ligand is much faster than the binding of a ligand from the bulk solution, a decomposition into the product of the survival probability of the receptor with only a single neighboring ligand and the survival probability of the receptor surrounded by an equilibrium distribution is possible and has been widely used (2, 24).

In a crowded environment, it is unclear whether, and under what circumstances, this timescale separation exists since diffusion of ligands to and away from the receptor become nontrivial and the distribution of ligands around the geminate pair may not be the equilibrium distribution. Hence, we aim to calculate 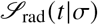 and 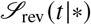 directly from simulation. From this, we can then determine the sensor autorcorrelation time and measurement error in the presence of crowding.

### Simulation Methodology

To obtain the correlation time from simulations, one possibility would be to directly model reversible reactions within a crowded environment. However, gathering robust statistics on survival times requires simulating the reaction-diffusion system for a period many times the correlation time of the receptor. Much of this simulation time is spent in the bound state, especially when the unbinding rate is large. During this time, the sensor is not gaining any additional information about the environment (7). Furthermore, as the volume fraction of crowders increases, the unbinding rate may decrease significantly (8), requiring even longer simulations. If the unbinding rate is slow compared to the timescale for the crowders to reach an equilibrium configuration around the bound complex, then the simulations can be significantly accelerated by decoupling the simulation of the bound and unbound states. This is valid when the relaxation time for the fluid around the bound complex, which is approximately captured by the molecular timescale 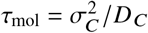, is small compared to the mean bound time,

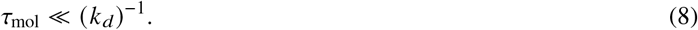

When this criteria is met, it is possible to calculate the probability of an initially bound complex being unbound at time *t* for a reversible first-order reaction in a crowded environment without explicitly simulating the reversible reactions.

#### Quasi-reversible particle-based reaction dynamics

In the simulation procedure proposed here, we effectively decouple the calculation of Eq. (7) into calculations of *k_d_* using simulations of the solution around the bound complex, and calculations of 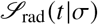 using simulation of the unbound receptor in solution. In particular, we decompose the system dynamics into diffusion in the bound state, unbinding, diffusion in the unbound state, and binding. Particle-based Brownian dynamics simulations (25) are used to obtain an equilibrium ensemble of configurations for the system when in the bound state. These configurations are then combined with Monte Carlo sampling of unbinding events to calculate the probability of unbinding in the crowded solution. This process also generates an ensemble of configurations of the system immediately following unbinding. Using these configurations as initial conditions, we then calculate 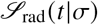.

Any simulation of reactions between physically interacting particles requires specification of a microscopic model for when reactions occur. Our general framework is independent of this microscopic model so to facilitate the description of the methodology, we will first assume a general reaction model, then specify the details of the reaction model used in our calculations in the following section. In general, a microscopic reaction model specifies the probability of transitioning from a state **x** in the phase space of the system with reactants to a state **y** in phase space with the product. If we let **x** represent the configuration of a system with *n_L_* ligands, *n_C_* crowders, and a single unbound receptor, and **y** represent the configuration of the system when one ligand has bound the receptor, then the microscopic association rate function is *k*_+_(**y**|**x**) and the microscopic dissociation rate function is *k*_+_(**x**|**y**).

Given a microscopic reaction model as specific through the rate functions *k*_+/−_, we calculate the separation probability in Eq. (7) in four steps: (i) Brownian dynamics simulation of the system in the bound state to generate an ensemble of configurations {**y**}, (ii) Monte Carlo sampling of unbinding transitions from {**y**} to states of the system immediately following an unbinding reaction {**x**}_0_, (iii) Brownian dynamics simulations with initial conditions specified by {**x**}_0_ to produce an ensemble of unbound trajectories {**x**(*t*)} in the unbound state, (iv) Monte Carlo sampling of along each trajectory to get an ensemble of reaction probabilities {*k*_+_(**x**(*t*))}, which can then be averaged to calculate the survival probability in the convolution in Eq. (7). We will now discuss each step in detail.

The first step involves a Brownian dynamics simulation of the system in the bound state. A single trajectory that is many times the length of the molecular timescale *τ*_mol_ provides a large number of independent samples from the equilibrium configuration surrounding the bound complex. In step (ii), this set of configurations {**y**} is used for Monte Carlo sampling of dissociation reactions. For each **y**_*i*_ ∈ {**y**}, a test dissociation move is proposed in which the products are placed at positions 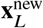 and 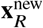, leading to a new configuration for the system in the unbound state **x**^new^. The specific choice of distribution from which new test positions for the products are drawn depends on the microscopic reaction model. The change of internal energy between the old and new state is computed, and the acceptance probability is determined by Metropolis–Hastings sampling. This probability, *p*_diss_, accounts for the excess contribution to the free energy of dissociation due to crowding. Importantly, we note that this is independent of the intrinsic dissociation rate in the absence of crowding, 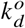 and the dissociation rate is given by

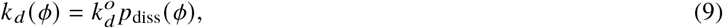

where *ϕ* denotes the dependence on crowding volume fraction. Since *p*_diss_ is independent of 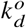, a single simulation provides *k_d_* for any intrinsic dissociation rate so long as the timescale separation Eq. (8) remains valid.

In addition to computing *p*_diss_, the configuration of each accepted trial reaction is saved to construct an ensemble of configurations of the system immediately following dissociation, {**x**}_0_. In Step (iii), each configuration is used as an initial condition for a Brownian dynamics simulation in the unbound state. In these simulations, the particles interact physically with one another, but do not react, leading to a trajectory **x**_*i*_ (*t_j_*) for each initial condition. For each trajectory *i* and time step *j*, the probability of an association reaction occurring is decomposed into the probability of a reaction being proposed based on an intrinsic association rate 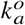, and, given a reaction is proposed, the probability it would occur based on the configuration of the system

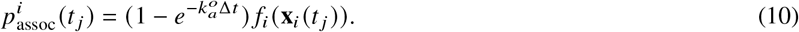

The first term gives the probability of a reaction occurring independent of the locations of particles. *f_i_*(**x**_*i*_ (*t_j_*)) is probability of a reaction occurring for the system in state **x**_*i*_, which depends on the microscopic reaction model and interaction energies between molecules. The survival probability of trajectory *i* is given by

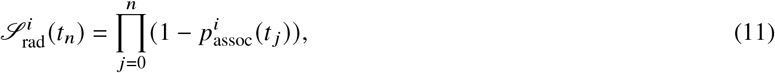

The survival probability is then found by taking the average over the ensemble of trajectories

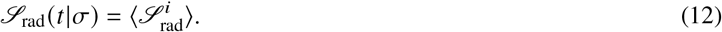

With the survival probability computed using Eq. (12), the autocorrelation function and the separation probability can be computed from Eq. (6) and Eq. (7), respectively. As with the case of the unbinding reactions, the calculation for *f_i_*(*t*) from the simulation data is independent of the intrinsic reaction rate 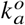 meaning that one set of Brownian dynamics simulations is sufficient to calculate the survival probabilities for any pair of intrinsic association and dissociation rate constants (under the constrain of Eq. (8) remaining valid). This provides significant gains in computational efficiency compared to fully reversible particle-based reaction-diffusion simulations, which require new simulations for each set of rate constants.

One limitation of the method as described is that each trajectory must be long enough for the survival probability to reach approximately zero. This limits the range of intrinsic association rates that can be used since small values of 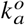 will lead to long survival times. However, the need for long simulations in the unbound state can be avoided by noticing that the system reaches a steady state over a timescale of approximately the molecular time *τ*_mol_. At times much greater than this, the reaction event density becomes constant, since the memory of the initial condition decays to zero and reactions are the same as between the receptor and the equilibrium distribution of ligands. The survival probability at these times can be readily extrapolated from the survival probability data as long as the data reaches this steady-state regime. Since the initial structure of the fluid decays relatively quickly, shorter simulations combined with extrapolations can produce survival distributions for small values of 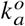 that would otherwise be computationally unfeasible. For the results presented in this work, we ensure simulations of the unbound state run well into the steady-state regime, then extrapolate the long-time survival probabilities through a least-squares fit of an exponential function to the survival probability data at late times (see Supplementary Material). This is equivalent to calculating the long-time limit to the time-dependent rate constant and using these values for reactions dynamics at long times.

#### Microscopic reaction model

While the general procedure described here can theoretically be used for any microscopic reaction model, we will specify our description to a recently-proposed reaction model for interacting particles, called interacting-particle reaction dynamics, that maintains detailed balance in equilibrium (26). This model captures the dependence of macroscopic reaction rates on crowding and provides a straightforward validation of our method. For a reversible bimolecular reaction, the transition rate functions are split into absolute proposal rate functions *λ*_+/-_, normalized proposal densities *q*_+/-_, and acceptance rates *α*_+/-_ such that

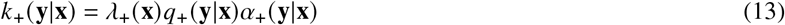

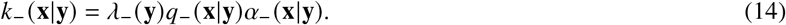

For the absolute proposal rate of association reactions, we use the Doi model (27) in which a ligand closer than a distance *R*_react_ has a constants proposal rate 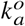 and a proposal rate of zero otherwise. The absolute proposal rate for dissociation reactions is a constant 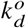, giving

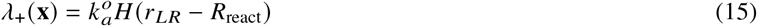

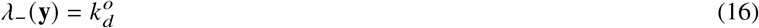

where *H*(*r*) is the Heaviside function and *r_LR_* = ║**x**_*L*_ – **x**_*R*_║ is the distance between the proposed reactants. The normalized proposal densities are chosen to maintain detailed balance by weighting the locations of the dissociation products according to the Boltzmann factor due to the interparticle interactions between the products. Doing so leads to (26)

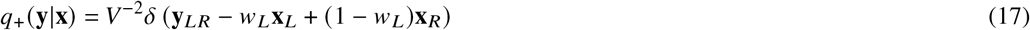

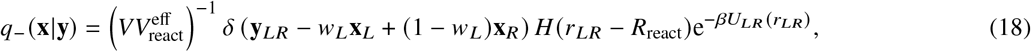

where *β* = (*k_R_T*)^−1^, *U_LR_*(*r*) is the potential energy between the products in the proposed dissociated configuration, *δ*(*x*) is the Dirac delta function which defines the placement of the product(s) of a reaction based on the locations the reactant(s) and a weight function *w_L_*. For all simulations in this work, we choose *w_L_* to be the ratio of the ligand volume to the that of the complex volume so that 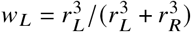, where *r_L_* and *r_R_* are the radii of the ligand and receptor, respectively. Hence, for equal sized ligands and receptors, the weights for both the ligand and receptor positions become equal. If *V* is the volume of the system, 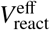 is an effective volume of the reactive space given by

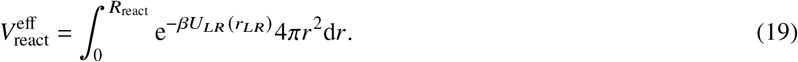

The acceptance probabilities account for the excess energy aside from the interactions between the products, which is accounted for in the probability densities of proposed locations. The association and dissociation acceptance probabilities are respectively given by

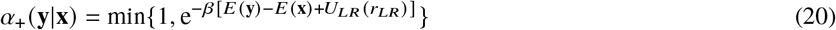

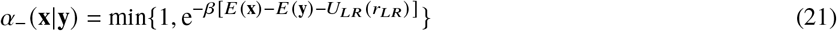

where *E*(**x**) and *E*(**y**) are the total internal energies of the system in the unbound and bound states, respectively.

For this microscopic reaction model, the excess association probability *f_i_*(*t_j_*) is calculated as follows. First, a candidate ligand for the reaction is chosen from the set of ligands that are less than a distance *R*_react_ from the receptor. If no ligands are within the reaction zone, *f_i_*(*t_j_*) = 0. Else, a proposed position of the product is drawn from the distribution give by Eq. (17), which is computed before the simulation and stored as a lookup table. *f_i_*(*t_i_*) is then given by the acceptance probability in Eq. (20).

### Validation of quasi-reversible simulations

To validate the proposed algorithm we compare the results from the quasi-reversible method with results from explicit simulations of the reversible reaction using the software package Readdy2 (25). We consider two scenarios for validation. In the first case the system consists of only a single ligand and single receptor in a periodic domain. This model can be compared to both the explicit reversible simulations and to known analytical results. In the second scenario, the system has two ligands, a receptor and an inert crowder molecule. This case provides a stringent test for the method in dense and crowded environments.

#### Single ligand, single receptor

The simplest validation of the proposed method is to consider a system of a single ligand-receptor pair that reversibly bind. The particles interact via a truncated Lenard–Jones potential with cutoff *r_c_* at the minimum of the potential, and shifted vertically such that the potential is purely repulsive and smooth. For *r* < *r_c_*, the potential between species *X* and *Y* is

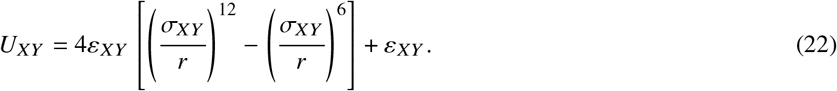

Throughout this work we set the length scale of interactions of a species with itself (e.g. *σ_X_*) and choose the interaction distance between two different species *X* and *Y* to follow the Lorentz-Berthelot rule, *σ_XY_* = (*σ_X_* + *σ_Y_*) /2. We set the units of length for the simulation by taking *σ_R_* = 1 and *σ_L_* = 1. Energy in the system in measured in units of *k_R_T* and *ε_LR_* = 0.667. The two particles diffuse freely with diffusion constants *D_L_* = *D_R_* = 1, which sets the timescale of the simulations to be *τ*_mol_ = 1, in a cubic simulation box of side length *I* = 2.5 equipped with periodic boundary conditions. The timestep used is Δ*t* = 10^−4^.

To compare the quasi- and fully-reversible simulations, we calculate the survival probabilities for different values of the intrinsic association rate constant as well as the mean and standard deviations of the receptor occupancy. Figure 1(a) shows the survival probabilities for explicit simulations and quasi-reversible simulations. The survival probabilities agree quite well for intrinsic association rates that span four orders of magnitude. Importantly, all survival curves from the quasi-reversible method are generated from the same set of binding and unbinding simulations, whereas each survival curve for the fully-reversible case requires an additional simulation. Figures 1(b)–(c) compares the mean receptor occupancy and variance of the occupancy for both simulation methods and the theoretical mean value. The theoretical mean values for the single reactive pair (26) are given by

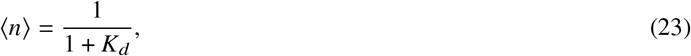

where

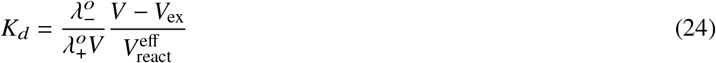

and

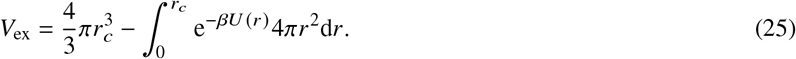

**Figure 1:**
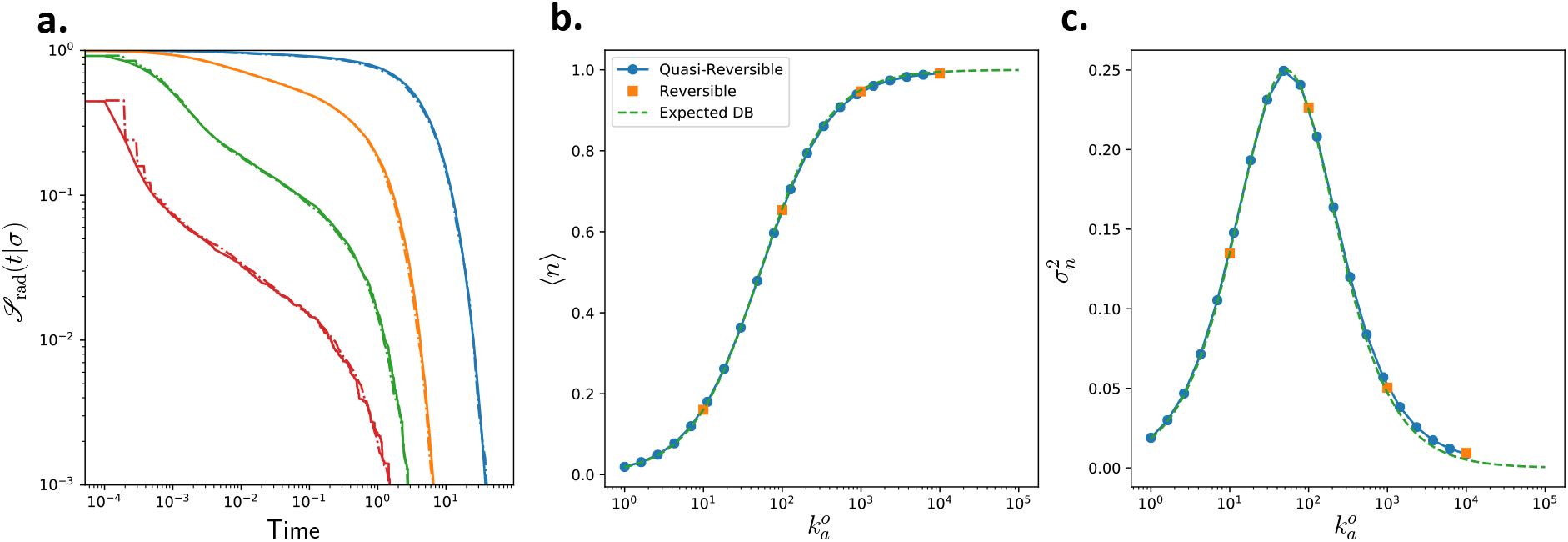
Validation of the quasi-reversible method for a system with a single ligand-receptor pair. Panel (a) shows the survival probabilities of an unbound receptor for different values of intrinsic association rate constant. Solid lines are from fully reversible simulations and dot-dashed lines are from a quasi-reversible simulation. Panels (b) and (c) show the mean occupancy and the variance in the occupancy of the sensor. For all simulations 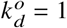.

The mean occupancy for the quasi-reversible simulations are computed as the ratio of the mean first passage times for unbinding and binding 〈*n*〉 = *τ_d_*/*τ_a_*, where

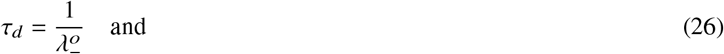

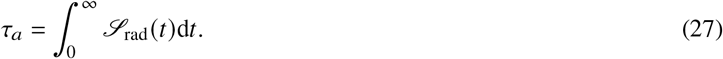

The result for the quasi-reversible simulations agree quite well with both the fully reversible interacting-particle reaction-diffusion simulations and the analytical result. As intrinsic association rate approaches the inverse of the timestep, both numerical methods differ from the analytical result since the approximation that the reaction probability in a given timestep is proportional to 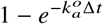 is no longer accurate.

#### Multiple ligands with inert crowder

To provide a more stringent validation test for the proposed method, we next consider a system that includes two ligands, one receptor and one inert crowder. In this case, there are four species that must be considered: ligands, the crowder, the unbound receptor and the ligand-receptor complex. The interactions between the species are chosen such that the ligands, crowders, and unbound receptors are of equal size, with *σ_L_* = *σ_C_* = *σ_R_* = 1, and the bound complex has a radius that preserves the volume of the binding ligand and receptor. Note that this does not mean that excluded volume remains the same upon binding. All interaction energies are set to be *ε*_LJ_ = 0.667*k_B_T*, and the diffusion constants are scaled according to particle radius such that *D_L_* = *D_C_* = *D_R_* = 1 and *D_LR_* = 2/*σ_LR_* so as to follow Stokes–Einstein scaling.

Figure 2(a) shows survival probabilities computed using fully-reversible simulations and the quasi-reversible simulation method. The two methods agree well for intrinsic rate constants that vary over five orders of magnitude and capture the full response range, as shown in Figure 2(b) (see Supplementary Material for a quantitative error analysis). The quasi-reversible method is also significantly faster than the fully reversible simulation. To compute the mean occupancy of the receptor to 0.1% accuracy for the case 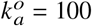, the quasi-reversible method requires only one-fifth of the computational time (47 vs 254 hours) of the reversible simulation method (see Supplementary Material for details). In addition to the greater efficiency for simulating a single parameter set, a clear advantage of the quasi-reversible method is that, once the reaction probabilities as a function of time have been computed once, quantities such as survival probabilities, mean occupancies or correlation functions can be computed for any combination of intrinsic rate constants at negligible computational cost since this becomes a simple post-processing step. Hence, the quasi-reversible method presented here can lead to significant speedup in the calculation of binding statistics for reversibly reacting systems. In the following section, we apply the quasi-reversible method to investigate the effects of crowding on the accuracy of a receptor-based sensing mechanism.

**Figure 2:**
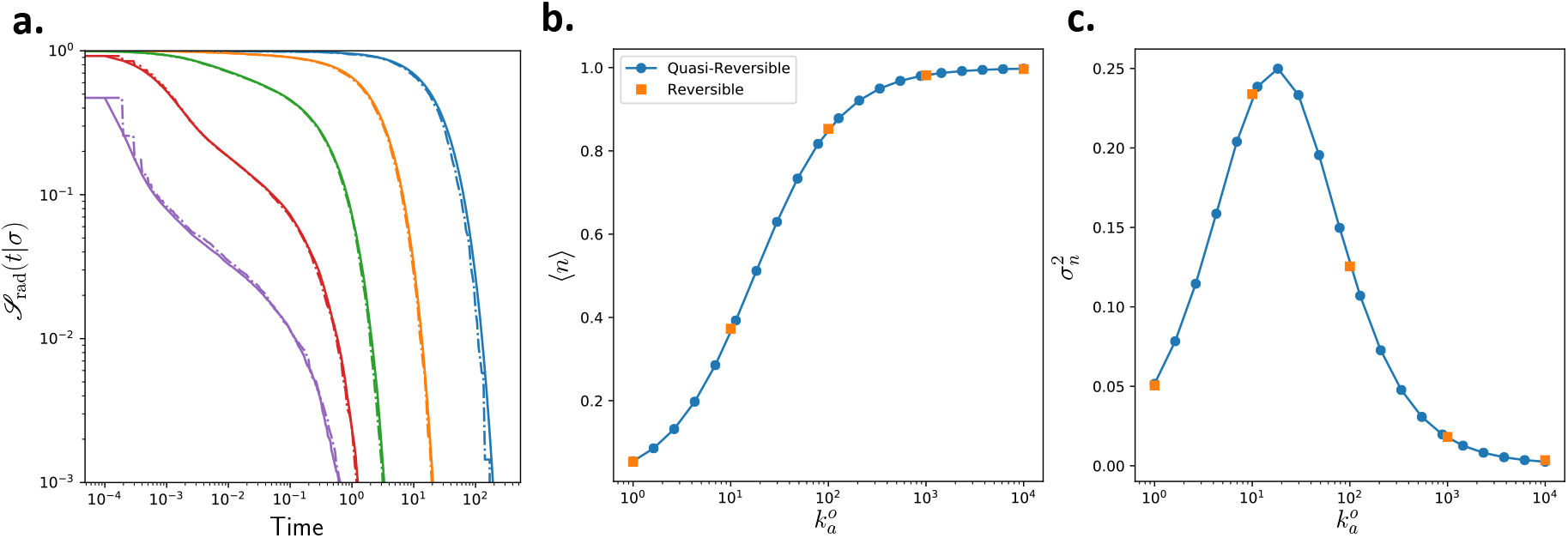
Validation of the quasi-reversible method for a system with a two ligands a receptor and an inert crowder. Panel (a) shows the survival probabilities for an unbound receptor to remain unbound for different values of intrinsic association rate constant. Solid lines are from fully reversible simulations and dot-dashed lines are from a quasi-reversible simulation. Panels (b) and (c) show the mean occupancy and the variance in the occupancy of the sensor. For all simulations 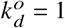. Quantitative comparison between the reversible and quasi-reversible simulations are presented the Supplementary Material.

## RESULTS

The intracellular environment is in constant flux. If cellular homeostasis is to be maintained in the face of external perturbations, it is crucial that cellular sensors of chemical ligands provide useful measurements even when macromolecular crowding is high. While much work has demonstrated how signal transduction networks in cells process and transmit information, it is often assumed that the ligands are non-interacting. In cells, however, ligands, especially when large proteins or RNA acts as the messenger, interactions with surrounding macromolecules and other ligands is the rule, not the exception. To better understand how crowding affects the accuracy of chemoreception inside of cells, we simulate two systems in which a receptor is surrounded by a mixture of ligands and inert macromolecular crowders. In one system, the ligands are of the same size as the receptor and the crowders, mimicking large messengers such as large proteins or RNA. In the other system, the ligand is smaller than the receptor and the crowders, as would be the case for small-molecule ligands, or small peptides. We assess how signaling in each of these systems is affected by macromolecular crowding.

The system consists of a single receptor, *n_L_* ligands and *n_C_* crowders in a cubic box of side length *I* and periodic boundary conditions. The particles interact via the cut and shifted Lennard Jones potential of the form in Eq. (22). The length scale for the simulations is set by the effective diameter of the receptor *σ_R_* = 1 and the timescale is set by molecular timescale of the receptor 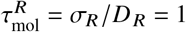, which fixes the diffusion constant of the receptor to be *D_R_* = 1. All parameters used in the simulations are given in Table 1.

**Table 1:** Simulation parameters.

| Parameter | Value |
| --- | --- |
| $n_L$ | 10 |
| $l$ | 5 |
| $\sigma_R$ | 1 |
| $\sigma_L$ | 1, 0.2 |
| $\sigma_{RL}$ | 1.26, 1.00266 |
| $\sigma_C$ | 1 |
| $\varepsilon_{LJ}$ | 1 |
| $\Delta t$ | $2 \times 10^{-5}$ |
| $N_{\text{traj}}$ | $> 10^4$ |
| $N_{\text{steps}}$ | $10^5$ |

<sup>a</sup> Parameters are in arbitrary units. Values for small-ligand simulations, where different, are shown in second column of values.

### How does crowding affect the survival time distribution of an unbound sensor?

The survival time distributions for recently-dissociated receptors show two dynamic regimes (Figure 3). Initially, there is a sharp decrease in the survival probability corresponding to rebinding of the recently dissociated ligand-receptor pair. The magnitude of this drop depends on the intrinsic association rate, which, when large leads to high probability that the dissociated ligand will rebind before escaping back into the bulk. When the intrinsic binding rate is high enough, this rebinding actually consists of two timescales (see Figure 3(f)). The first, very short timescale, involves the recently dissociated molecule rebinding before ever leaving the reaction zone. For all but the largest values of 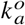, the probability of reacting over the course of this timescale is negligible. This is followed by a range with a less steep decrease in survival probability due to the recently unbound ligand diffusing a short distance away from the receptor before returning and rebinding. The rebinding probability, measured by the magnitude of the initial decrease, is proportional to the degree of crowding for the small-ligand case. This is due to the surrounding molecules acting as a diffusional barrier for the recently dissociated ligand, effectively corralling it in the vicinity of the receptor and increasing the number of encounters before escaping. For larger ligands, the rebinding probability is relatively insensitive to the degree of crowding.

**Figure 3:**
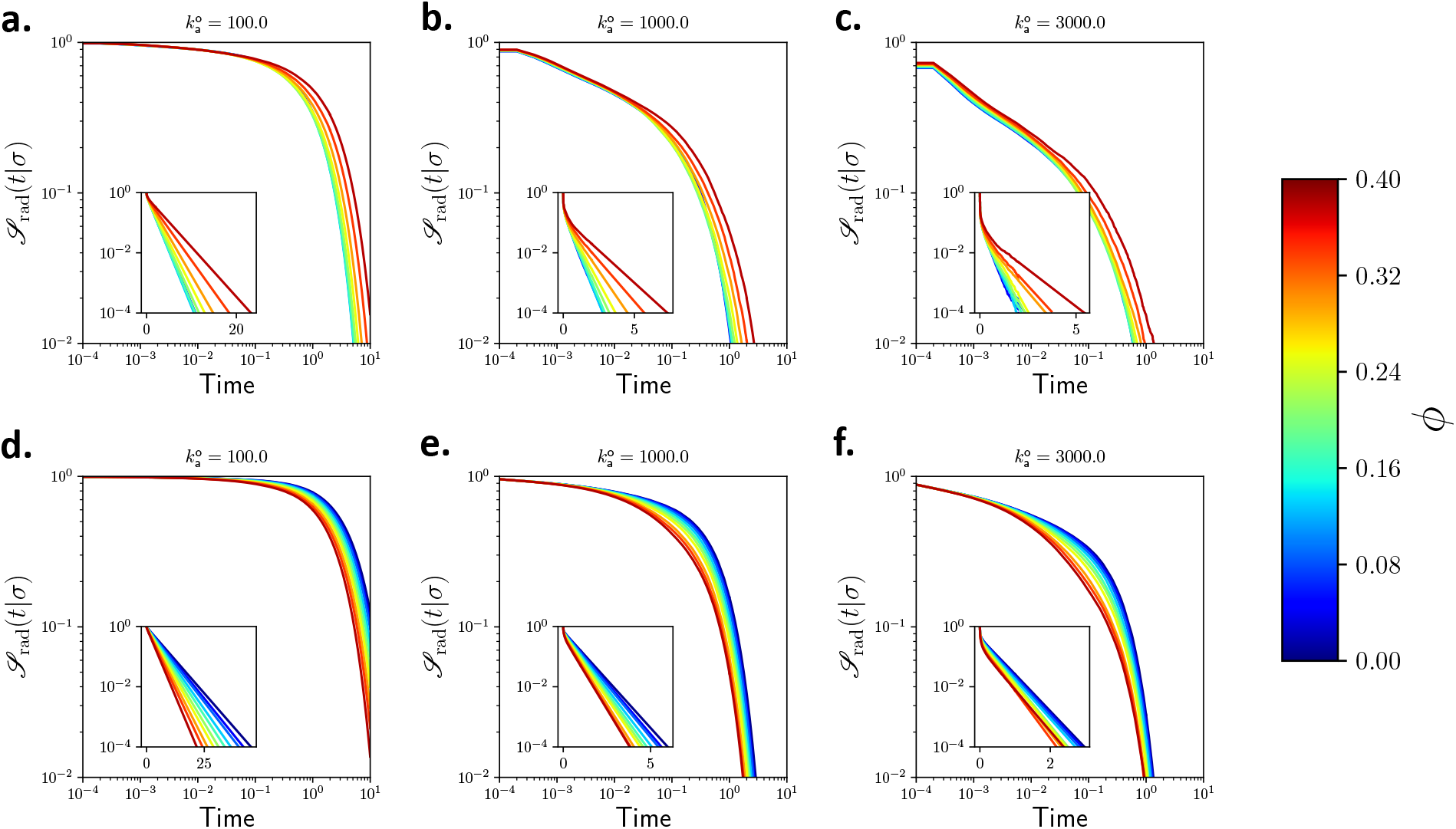
Survival probabilities for receptors in crowded solutions. Top row shows results for a ligand with radius *σ_L_* = 0.2. Bottom row shows results for ligand with radius *σ_L_* = 1. Each column corresponds to a different intrinsic rate constant: (left) 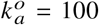, (b) 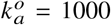, and (c) 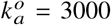. Insets show same data on a semi-log plot, with the long-time limit of the association rate being proportional to the slope. In all cases, *σ_R_* = *σ_C_* = 1, 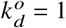, *N*_traj_ = 2 × 10^4^.

At longer times (Figure 3 insets), the survival probability is governed by ligands from the bulk solution encountering the receptor. Over these times, the survival probability decreases exponentially with a well-defined (long-time) rate constant. The long-time exponential decrease in survival probability shows the opposite dependence on crowding for the small and large ligand cases. In the case of small ligands, the long-time reaction rate increases with crowding. The change with crowding is greatest for small intrinsic rate constants. When the intrinsic rate of association is small, many contacts are needed on average for a reaction to occur. Crowding around the receptor corrals ligands in the vicinity of the receptor, facilitating a greater number of contacts before the ligand escapes back into the bulk.

In the case of larger ligands, the long-time limit of the time-dependent reaction rate shows the opposite trend with crowding. In this case, crowding slows the reaction. This could be due to either a decrease in the encounter rate as diffusion is slowed by the presence of more crowders, or the result of crowding making the conformational change from the encounter complex to the bound complex less favorable. To distinguish between these two effects, we calculate the average acceptance probability of association and the average encounter probability as a function of crowding. These correspond to calculating the mean value of *α*_+_(**y**|**x**) and *H*(*r_LR_* – *R*_react_), respectively. We calculate the average values over the set of trajectories and at long times, denoted by 〈〉_eq_, so that they correspond to the long-time limit of the reaction rates. Comparing Figure 4 (a) and (b), we see that crowding has a much larger effect on the transition energy than on the encounter rate, indicating that the slow-down observed with crowding is the primarily the result of the conformation change upon binding. For the small ligand, this effect is much less significant since the volume change of the bound and unbound receptor is much smaller.

**Figure 4:**
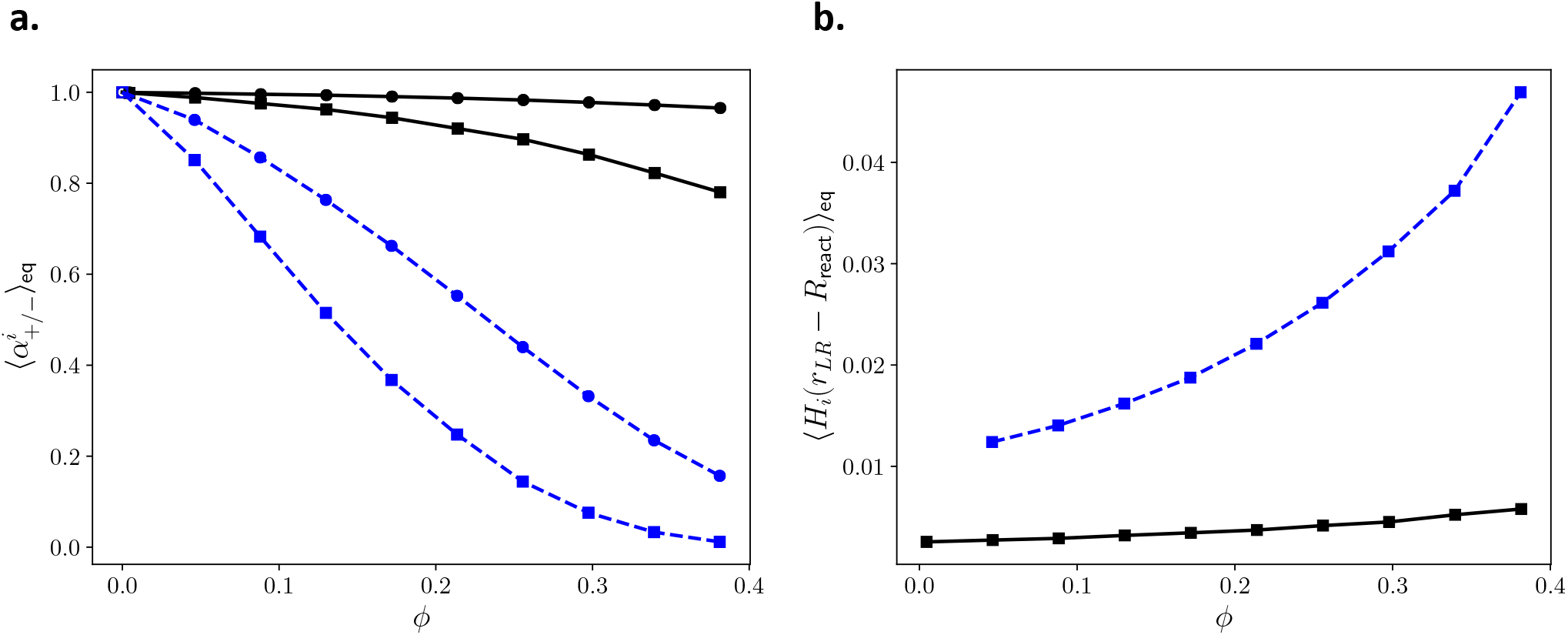
Panel (a) shows the excess transition probability due to crowding for association (circles) and dissociation (squares) reactions for ligands with *σ_L_* = 1 (dashed blue lines) and *σ_L_* = 0.2 (solid black lines). The unfilled points for zero volume fraction are the theoretical values for a dilute systems. Panel (b) shows the probability that any ligand is at a distance *r* < *R*_react_ at a given time. Blue dashed lines are for cases with *σ_L_* = 1 and black solid lines are for *σ_L_* = 0.2.

### How do the intrinsic reaction rates change with crowding?

In general, crowding alters the thermodynamics of both binding and unbinding due to differing interactions of the complex and the reactants with the surrounding solution. The change in reaction rates depend on the change in excess chemical potential between the bound and unbound states, which for simple fluids depends on the relative shapes and sizes of the bound and unbound states. For our microscopic model, the decrease in 〈*α^i^*〉_eq_ for both association and dissociation as shown in Figure 4(a) means that the microscopic association and dissociation rates both decrease as the degree of crowding increases. This is because, as crowding increases, it becomes more likely that a proposed reaction product (either the complex or the dissociated ligand-receptor pair) will overlap with nearby crowding molecules, making the transition less energetically favorable.

For the larger ligand, this effect is much more pronounced in both association and dissociation reactions. For association, the complex forms a single spherical particle whose volume equals that of the two reactants, and this is prevented by surrounding crowding molecules. Physically, this corresponds to a conformational rearrangement of the encounter complex into the bound complex, which is hindered in crowded solution. Dissociation also requires a cavity in the solution that allows the placement of the products back into the encounter complex conformation, and is also hindered by crowding. While these effects are still present in the small-ligand case, they are much less pronounced. In particular, the conformational change of the ligand and receptor into the bound complex is quite small, and is hence essentially unaffected by the crowded solution. The dependence of microscopic rate constants on crowding is specific to the chosen representation of the transition state complex and bound pair. For example, should both the transition state and bound complex be modeled with the same shape, which would be a valid approximation of some transient protein-protein interactions, there would be no change in the intrinsic rates due to crowding. For our volume-conserving model, the change is both in shape and in excluded volume, and as can be seen in Figure 4(a), it can have a substantial impact on the microscopic rates.

While increased crowding decreases the probability of an encountered ligand-receptor pair crossing the energy barrier to form the bound complex, it also alters the stead-steady distribution of ligands around the receptor. Figure 4(b) shows the probability that a ligand is within the reactive zone (i.e. *r_RL_* < *R*_react_) as a function of crowding. For both small and large ligands, crowding increases the likelihood that a ligand is in contact with the receptor. The important comparison is between this increase and the corresponding decrease in the probability of crossing the transition barrier from the encounter complex to the bound complex. If the local ligand enrichment is large enough, it will overcome the increased excess free energy of the binding transition, and the overall rate of ligand-receptor binding will increase, even though the microscopic transition rate decreases. When the excess free energy of binding is larger, the ligand-receptor association reaction rates will decrease.

Since both the microscopic association rate and dissociation rate are decreased by crowding molecules, it seems plausible that the correlation time in the receptor dynamics will increase on account of a general slowing-down of the reactions. However, the quantity of interest for the accuracy of a receptor’s predictions of the surrounding concentrations is the time between binding events of different ligands. If the same ligand continually rebinds the receptor, no new information about the concentration is being gleaned. Hence, it is plausible that there may be scenarios in which decreasing the association rate allows recently unbound ligands to escape back into the bulk and a new informative ligand can bind the receptor.

### Does crowding decrease the accuracy of a chemoreceptor?

The relative error of a concentration measurement is directly proportional to the correlation time of the receptor occupancy (see Eq. 2). Hence smaller correlation times correspond to more accurate sensing. Figure 5 shows the dependence of the receptor correlation time on the degree of crowding for a range of intrinsic association rate constants. For ligands of the same size as the receptor and crowder (Figure 5(a)) the correlation time monotonically increases with crowder volume fraction. This suggests that crowding degrades sensor accuracy when the ligand, receptor and crowders are of the same size, at least for the volume conserving microscopic model employed in this work. For the small-ligand case on the other hand, the correlation time can decrease with increasing crowding for high intrinsic association rates. How does crowding decrease the correlation time? To answer this question, we consider effective rates of binding and unbinding that coarse-grain out high-frequency, uninformative rebinding events.

**Figure 5:**
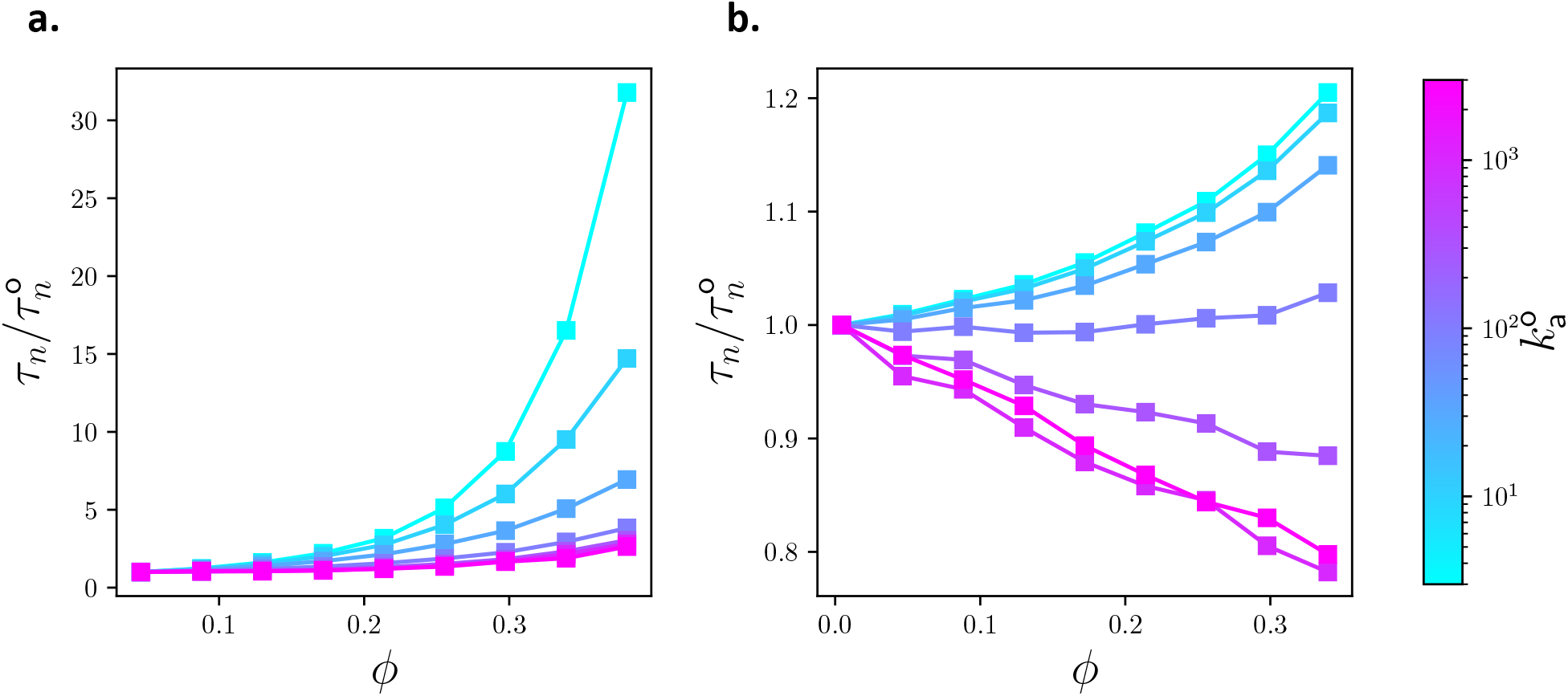
Correlation time of receptor as a function of crowding. Left panel shows correlation times normalized by the *n_C_* = 0 case for the large ligand (*σ_L_* = 1) case. Right panel is for the small ligand (*σ_L_* = 0.2) case. The intrinsic dissociation rate for all curves is 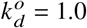.

In a diffuse system, it is possible to coarse-grain the sensor dynamics by integrating out the high frequency rebinding reactions that provide no information about the ligand concentration (1, 2). The end result is a two-state Markov model with effective switching rates that correspond to information-containing binding events and has the same correlation time as the fine-grained model. The effective on- and off-rates of (uncorrelated) ligand-receptor binding are related to the mean and correlation time of the fine grain simulation through

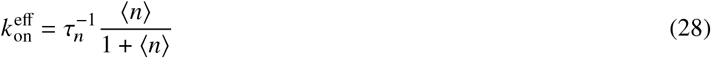

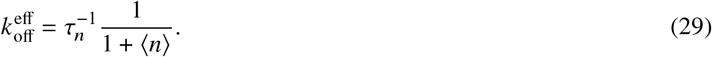

Figure 6 shows the dependence of the effective rates on crowding for a range of intrinsic association rates. In order for a new ligand to bind to an initially bound receptor two things must happen. First, the bound ligand must unbind and escape fromthe receptor back into the bulk solution without rebinding. Second, another ligand must approach the receptor from the bulk solution and bind to the unoccupied receptor. The effective off-rate measures the rate at which bound ligands unbind and escape, while the effective on-rate measures the rate at which unbound ligands arrive at that receptor and bind. Both effective rates are therefore functions of the diffusion rate of ligands and the intrinsic reaction rate once the encounter complex has formed.

**Figure 6:**
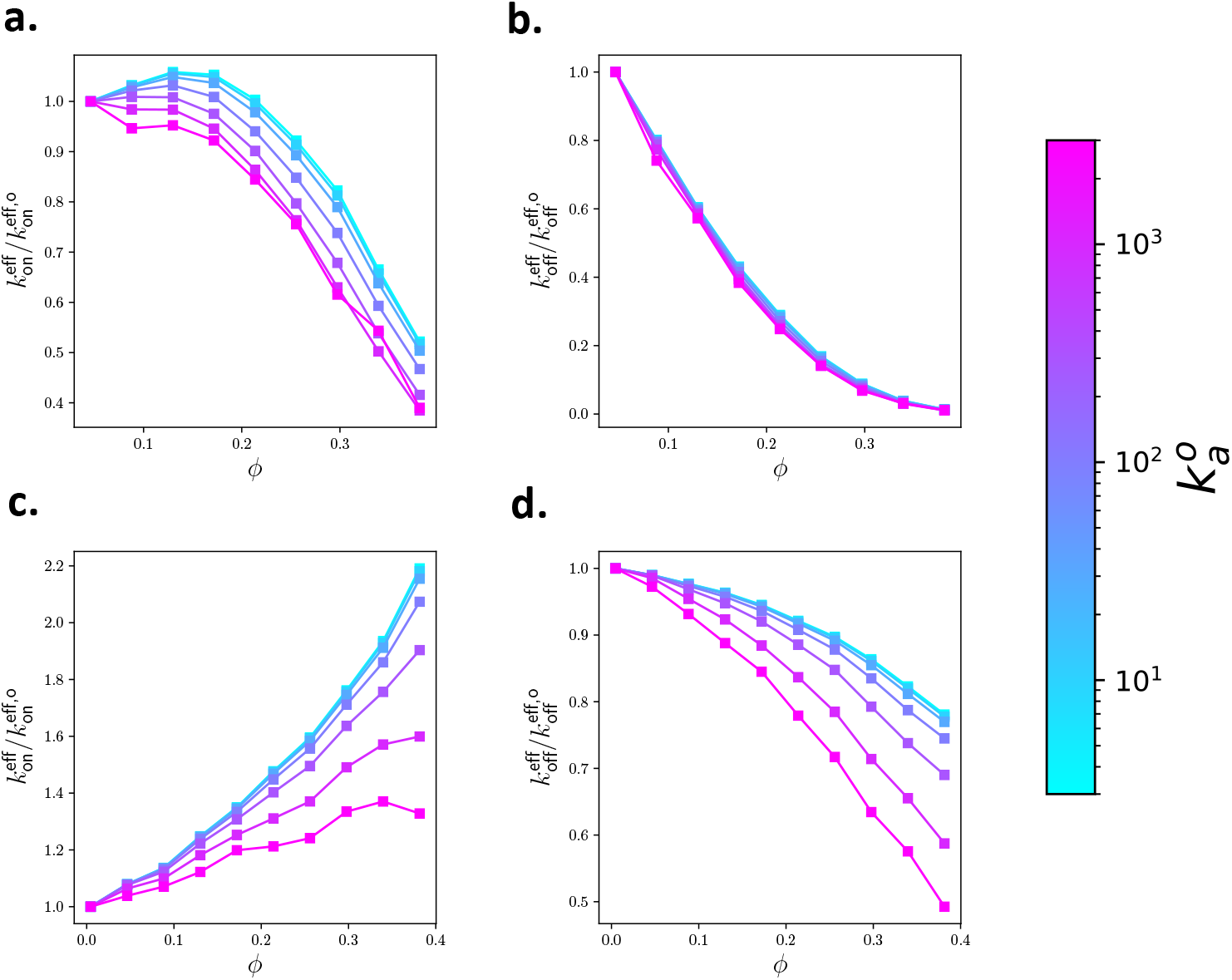
Effective switching rates for the coarse-grained sensor model. Top two panes show effective on (left) and off (right) rates for large-ligand case with *σ_L_* = 1. Bottom two panels show effective rates for small-ligand case with *σ_L_* = 0.2. The intrinsic dissociation rate for all curves is 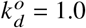.

For the large-ligand case, the effective on-rate is non-monotonic in the volume fraction of crowding for low intrinsic rate constants (Figure 6(a)). This is in agreement with previous studies of irreversible reacting hard-spheres (16). At low volume fractions, the effective on-rate increases because the volume occupied by inert crowding molecules increases the effective concentration of the ligands making them more likely to bind the receptor, but also only weakly decreases the diffusion rate of the ligands. As the crowder volume fraction increases, the reaction rate decreases due to the lower flux of ligands to the receptor through the crowded media. The dependence of the on-rate on crowding is also a function of the intrinsic association rate (16). When the association rate is high, ligands that encounter the receptor react with high probability so that only transient contact is required. For low intrinsic reaction rates, many more encounters (or time spent in the reaction volume) are needed for a reaction to occur. Hence these unlikely reactions are actually made more probable by small amounts of crowding since this slows the escape of ligands away from the reaction volume while not prohibiting the approach of new ligands too severely. This effect is more strongly present when the ligands are small Figure 6(c). In this case, the effective on-rate continues to increase with increasing crowding, and the increase is greatest for reactions with small intrinsic association rates.

The effective off-rate, shown in Figure 6(b) and (d), incorporate the intrinsic rate of dissociation and the time required for an unbound ligand to diffuse away from the receptor. 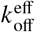 decreases monotonically with both crowding volume fraction and intrinsic association rate. Crowding affects the effective off-rate in two ways. It first decreases the intrinsic dissociation rate as seen in Figure 4(a). Second, it decreases the rate at which the dissociated ligand diffuses away from the receptor. This second effect is sensitive to 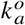 since an increased intrinsic association rate increases the likelihood of rebinding. By comparing panel (b) and (d) in Figure 6, we see that the effective off-rate of the larger ligand is dominated by the decreased microscopic dissociation rate, since it depends only very mildly on the 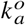. In contrast, the smaller ligand is more heavily influenced by the diffusion and caging effects of crowding, since these depend on the intrinsic association rate. The correlation time, which is inversely proportional to both effective on- and off-rates, decreases when the increase in effective on-rate is greater than the decrease in off-rate. For the small ligand, this occurs when the intrinsic association rate is high.

Crowding can lengthen the time a receptor and ligand remain correlated by more than an order of magnitude. This timescale, 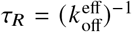, is called the residence time and is of particular interest to pharmacology and drug design (12). Figure 6(b) and (d) demonstrates that this timescale is not only a function of the dissociation rate, but depends sensitively on both the association rate and on the environment in which the reaction is taking place.

## DISCUSSION

In this work, we determined how the sensing accuracy of a receptor that reversibly binds ligands is impacted by macromolecular crowding. The autocorrelation time of the receptor occupancy, and hence the relative error of a concentration inference based on the receptor’s occupancy, can decrease with crowding under certain circumstances. In particular, when crowding facilitates binding of the ligand to the receptor, while also allowing rapid exchange of ligands with the bulk solution. In our simulations, we demonstrated that this is the case for a small ligand with a large intrinsic association rate. This indicates that crowding has the potential to increase the accuracy with which receptors can measure their chemical environment. The effective reaction rates provide intuition into why this is the case. To make an independent concentration measurement, a previously-unbound ligand must react with the receptor. The time between such events sets the rate at which the error in concentration measurement declines. This time depends on both the approach of a ligand to the receptor and the escape of a previously bound ligand from the vicinity of the receptor. If the increase in effective on-rate outweighs the decrease in effective off-rate, the correlation time of the sensor occupancy can decrease, allowing the sensor to make more independent measurements in the same amount of time. In particular, our simulations demonstrate this is feasible for small signaling molecules, such as short peptides or ion.

Examining the change in microscopic rate constants as a function of crowding demonstrates how important the microscopic model can be to understanding long-time dynamics of crowded systems. In our model, the microscopic rates are thermodynamic quantities related to the excess chemical potentials of the transition state complex and the bound complex. The sensitivity of the rate constants to crowding is the result of conformational changes upon binding that require a reorganization of the surrounding solution. These terms would likely be significantly smaller if the bound complex was a tangent dimer, especially when the signaling ligand is large, for example if a protein is the signaling molecule. Alternatively, if a significant conformation change is required to reach the bound state from the transition complex, such as the opening of a binding pocket, the effect of crowding could be even more significant than in our volume-conserving model. While changes to the microscopic rate constants clearly alter the high-frequency dynamics of the receptor state, they also impact the low-frequency switching of uncorrelated binding events. This is due to the impact that fast rebinding has on the escape probability of a nearby ligand. Our simulations show that crowding increases the likelihood of rebinding events, which in turn increases the correlation time of the sensor state, providing less accurate measurement of the surrounding environment.

Our analysis also demonstrates how the residence time of a ligand-receptor complex is fundamentally linked to the intrinsic association rate and diffusion. Furthermore, these quantities, along with the intrinsic dissociation rate depend quite heavily on the degree of macromolecular crowding. This could have important implication for drug development, where the residence time is an important metric for efficacy (12). Our simulations show two important considerations. First, the residence time as measured though an in vitro dissociation assay may not provide a meaningful measure of the residence time inside of a cell. Second, drugs that show the same dissociation rate outside of a cell may display different dynamics inside a cell depending on the mechanism through which crowding impacts the effective dissociation rate. Our results should motivate experimental pharmacokineticists to develop models of the crowded cellular environment to understand systematically how residence time can change inside cells.

In addition to investigating the effect of macromolecular crowding on the accuracy of cellular chemoreception, we have developed a novel algorithm for generating robust statistics of reversible reactions between interacting particles in dense systems. The method developed here generates survival probabilities of both the bound and unbound states by considering reaction probabilities along an ensemble of independent trajectories. The method takes advantage of the timescale separation present in many biochemical systems between the time a ligand-receptor pair remain bound and the time the surrounding solution relaxes to the equilibrium distribution. By decoupling the binding and unbinding reactions, one set of simulation trajectories produces survival distributions and separation distributions for a broad range of intrinsic association and dissociation rates. This greatly reduces the computational cost for evaluating the impact of the microscopic model on low-frequency reversible reaction dynamics. In the future, this method could readily be used in conjunction with optimization methods to find a microscopic model that minimizes a desired cost, which otherwise might be unfeasible if individual simulations are required for each cost-function evaluation. For example, one could determine the intrinsic rate constants that maximize the mutual information for a signaling reaction in a crowded environment.

## CONCLUSION

In this work we have used a novel simulation method to model the binding kinetics of a ligand-receptor system in a crowded environment. Our results show that the accuracy with which the receptor occupancy can infer the concentration of ligands can increase with greater macromolecular crowding. Whether the accuracy degrades or improves with crowding depends nontrivially on the size of the ligand, and on how the intrinsic association and dissociation rate constants, and the diffusion rate change with increased crowder volume fraction. Additionally, by determining effective rate constants that integrate out rapid rebinding events between the ligand and receptor, we show that the residence time of a ligand can be very sensitive to macromolecular crowding. This sensitivity arises from both thermodynamic alterations to the microscopic binding process and dynamical effects resulting from the escape probability of a ligand in the vicinity of a receptor. Lastly, the simulation method presented here can readily be applied to other problems in which reversible biomolecular reactions under physiological conditions are of interest.

## AUTHOR CONTRIBUTIONS

W.S. and S.S. designed the research. W.S. carried out all simulations, analyzed the data. W.S. and S.S. interpreted the results. W.S. and S.S. wrote the manuscript. All authors gave approval of the final version of the manuscript.

## ACKNOWLEDGMENTS

This work is partially supported by the University of Michigan Protein Folding Diseases Initiative. W.S. was partially funded through the Michigan IRACDA Program K12 GM111725.

## SUPPLEMENTARY MATERIAL

Supplementary Material containing 5 figures and 5 equations is available online. All codes used to generate and analyse results are available at https://github.com/stroberg-lab/qrevIPRD.

